# Protection against avian coronavirus conferred by oral vaccination with live bacteria secreting LTB-fused viral proteins

**DOI:** 10.1101/2021.10.04.462992

**Authors:** Avishai Lublin, Chen Katz, Nady Gruzdev, Itamar Yadid, Itai Bloch, Yigal Farnoushi, Luba Simanov, Asaf Berkovitz, Dalia Elyahu, Jacob Pitcovskib, Ehud Shahar

**Affiliations:** The Department of Avian Diseases, Kimron Veterinary Institute, Israel; MIGAL Research Institute in the Galilee, Kiryat Shmona, Israel; Tel-Hai Academic College, Upper Galilee, Israel

**Author notes:** Equal contribution (Avishai Lublin, Chen Katz).

**Keywords:** Infectious bronchitis virus, Avian corona virus, Oral vaccination, Heat labile enterotoxin, Subunit vaccine, Fusion polypeptides

## Abstract

The devastating impact of infectious bronchitis (IB) triggered by the IB virus (IBV), on poultry farms is generally curbed by livestock vaccination with live attenuated or inactivated vaccines. Yet, this approach is challenged by continuously emerging variants and by time limitations of vaccine preparation techniques. This work describes the design and evaluation of an anti-IBV vaccine comprised of *E. coli* expressing and secreting viral spike 1 subunit (S1) and nucleocapsid N-terminus and C-terminus polypeptides fused to heat-labile enterotoxin B (LTB) (LS1, LNN, LNC, respectively). Following chicken oral vaccination, anti-IBV IgY levels and cellular-mediated immunity as well as protection against virulent IBV challenge, were evaluated 14 days following the booster dose. Oral vaccination induced IgY levels that exceeded those measured following vaccination with each component separately. Following exposure to inactivated IBV, splenocytes isolated from chicks orally vaccinated with LNN or LNC -expressing bacteria, showed a higher percentage of CD8^+^ cells as compared to splenocytes isolated from chicks vaccinated with wild type or LTB-secreting *E. coli* and to chicks subcutaneously vaccinated. Significant reduction in viral load and percent of shedders in the vaccinated chicks was evident starting 3 days following challenge with 10^7.5^ EID_50_/ml virulent IBV. Taken together, orally delivered LTB-fused IBV polypeptide-expressing bacteria induced virus-specific IgY antibody production and was associated with significantly shorter viral shedding on challenge with a live IBV. The proposed vaccine design and delivery route promise an effective and rapidly adaptable means of protecting poultry farms from devastating IB outbreaks.

**Highlights:**

- Mucosal vaccination was shown particularly beneficial against respiratory viruses.
- An anti-IBV vaccine composed of three IBV polypeptides fused to LTB was designed.
- Vaccine composed of bacteria secreting polypeptides was orally delivered.
- Vaccine induced specific immune responses and shortened viral shedding duration.

## Introduction

Infectious bronchitis (IB) is a highly contagious acute respiratory disease, primarily affecting chickens, worldwide [1]. Other non-respiratory clinical manifestations of IB are nephritis, and reduced egg production and quality. IB virus (IBV), the etiological viral agent of the disease, is an enveloped *Coronaviridae*, of the genus *Gammacoronavirus*, which is grouped into six, genetically divergent genotypes (GI-GVI) [2], that all contain a non-segmented, single-stranded, positive-sense, ~27,600-nucleotide RNA genome [3]. The IBV genome encodes 15 non-structural proteins involved in replication (ORF1ab), four accessory proteins and four structural proteins (spike glycoprotein [S], matrix [M], nucleocapsid [N], and envelope protein [E]) [4, 5]. The S protein is comprised of two subunits, with S1 mediating viral attachment to host cells and bearing a receptor-binding domain (RBD), and S2 driving membrane fusion [6]. Antigenic and genetic variants of IBV have been classified based on S1 gene nucleotide sequence analysis.

IBV infects the host by inhalation or ingestion and initially replicates in the upper respiratory tract and trachea, the primary target organ, after which, it disseminates to enteric tissues, particularly the proventriculus, intestine and cecal tonsils [7], as well as to the kidneys [8] and the oviduct [9]. Infection of the lymphoid tissue-containing cecal tonsils in the intestine commonly results in a persistent infection lasting weeks or months after the acute disease episode [10, 11]. IBV can rapidly spread, and cause high rates of morbidity and mortality [12].

The most efficient means of coping with IBV is via vaccination with either live attenuated or inactivated vaccines. The recommendation by veterinary authorities in Israel for broilers is to administer an attenuated vaccine at 1-day of age by spray, drinking water or eye drop, followed by a boost at 10-12 days of age. In layers and breeders, vaccination programs include live vaccines at young ages, followed by injection of inactivated vaccines. However, available vaccines are challenged by the emergence of new IBV serotypes, for which the vaccine strain provides little or no cross-protection [13].

Development of new live attenuated vaccines is performed by serial passage of field virus isolates in embryonated eggs, which is a laborious and time-consuming process with an unpredictable outcome concerning vaccine safety and degree of attenuation [14]. In the era of molecular biology, protein subunits for vaccination may be delivered as polypeptides, by viral or bacterial delivery systems that carry and express relevant genes encoding desired antigens [15–20]. For example, oral administration of *Lactococcus lactis* expressing MSA2, a protein projected to protect against *Plasmodium falciparum* infection, resulted in high IgG levels in vaccinated rabbits [21].

Adjuvants are often used to enhance immune responses to vaccination. One such mucosal adjuvant is heat labile enterotoxin B (LTB), which is a non-toxic homo-pentamer subunit of LT toxin expressed by enterotoxigenic *E. coli* strains (ETEC), responsible for binding to the host GM1 ganglioside receptors. LTB has been used in several vaccine development studies both as a free adjuvant and in fusion with various antigens. Subcutaneous immunization of mice with *H. pylori* urease antigen mixed with LTB, increased levels of specific IgG in serum and IgA in saliva and conferred high protection against the pathogen as compared to none-adjuvanted immunization and the untreated control groups [22]. Similarly, addition of LTB to egg drop syndrome adenovirus rKnob significantly elevated the antibody response of orally and transcutaneously vaccinated chickens [23]. Oral immunization of BALB/c mice with a chimeric protein of LTB and two epitopes from the Zaire ebolavirus GP1 protein, induced both IgA and IgG antibody responses [24].

The present work aimed to develop a trivalent anti-IBV vaccine comprised of LTB-fused viral S1 and two subunits of N proteins expressed and delivered by a live bacterial system. The humoral and cell-mediated immune responses, as well as, the protection efficacy of the oral vaccination following challenge were assessed in chickens.

## Materials and Method

### Cloning and expression of LTB-fused N and S1 IBV polypeptides

#### Generating structural models of LTB and viral antigens

The design of the fusion polypeptides with proper folding and correct epitope presentation was supported by a three-dimensional computer model for each of the fusion constructs. The following PDB codes were used as templates for homology modelling: *Carrier*: heat-labile enterotoxin B chain (LTB) – PDB IDs: 1EFI, 1FD7 and 1LTB; *Nucleocapsid protein* N terminal segment (amino acids 29-160) and C terminal segment (amino acids 218-326): 2BTL, 2BXX, 2GEC, 2CA1, 2GE7 and 2GE8; *Spike1 glycoprotein* (S1): 6CV0. Each template of a fusion protein component was manually oriented to allow pentameric complex formation for the LTB segment and presentation of the N and S1 epitopes. Thereafter, the Modeller program [25] was used to generate homology models of the fused protein, allowing for loop optimization in areas without an appropriate template structure.

#### Plasmid construction and transformation

DNA fragments coding for LTB and specific IBV domains, were ordered from GenScript. The oligonucleotide primers used to amplify the codon-optimized sequences are listed in Table 1. Briefly, each sequence was amplified in two steps, using primers F1 and F2 for all samples and R, Ra, Rb and Rc for LTB, LTB-NN (LNN), LTB-NC (LNC) and LTB-S1 (LS1), respectively. The resulting cassettes were cloned into the pGEM-T-Easy Vector (Promega), according to the manufacturer’s instructions. A schematic model of the constructs is presented in Figure 1. The plasmid constructs were transformed into electrocompetent *E. coli* MG1655 cells, using established procedures (Sambrook et al., 1989). Following transformation, several colonies growing on LB agar plates supplemented with 100 μg/ml ampicillin, were tested for the presence of the correct plasmid by colony polymerase chain reaction (PCR) with M13F and M13R primers and then by sequencing.

**Table 1.**
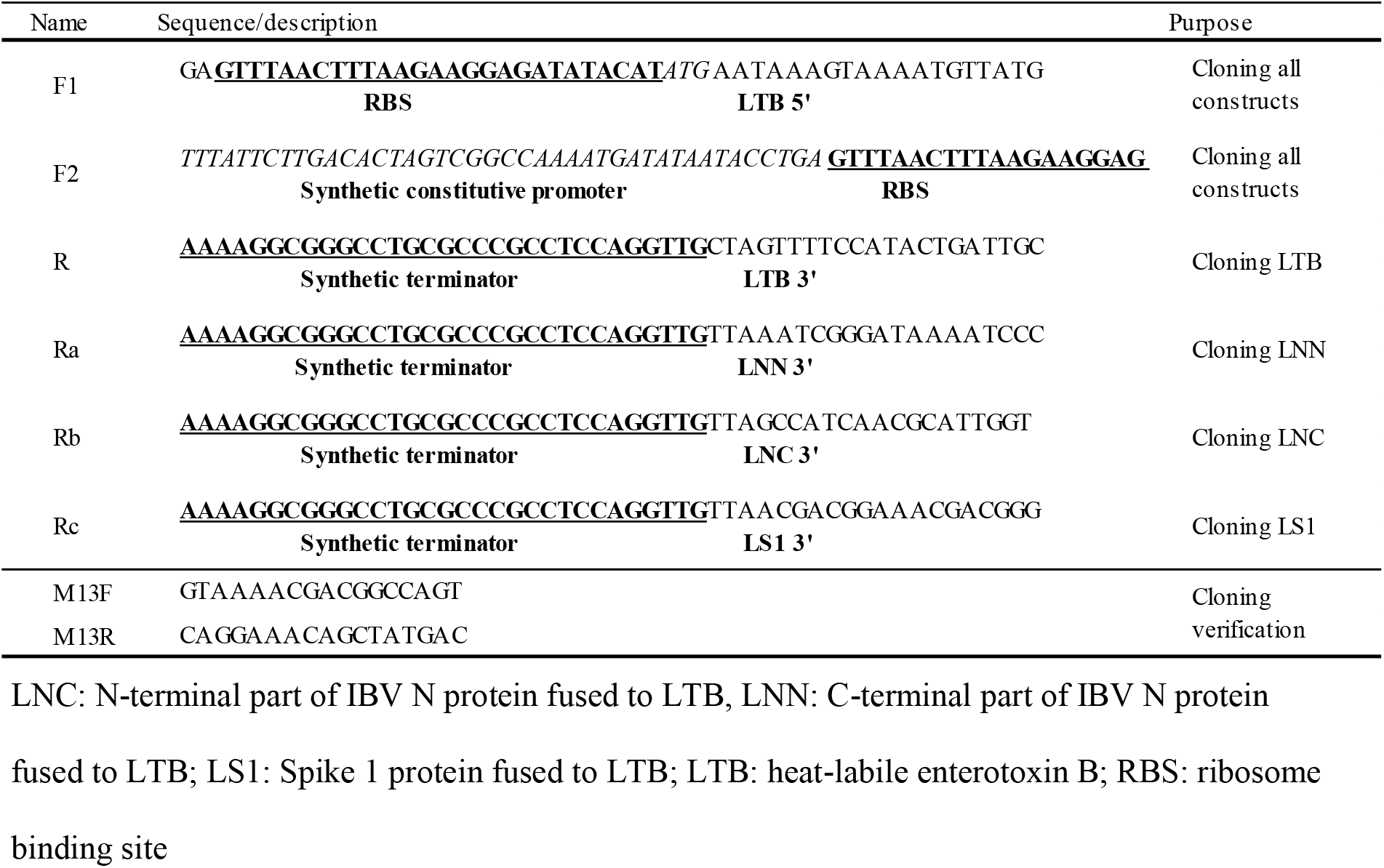
Primers list.

**Figure 1.**
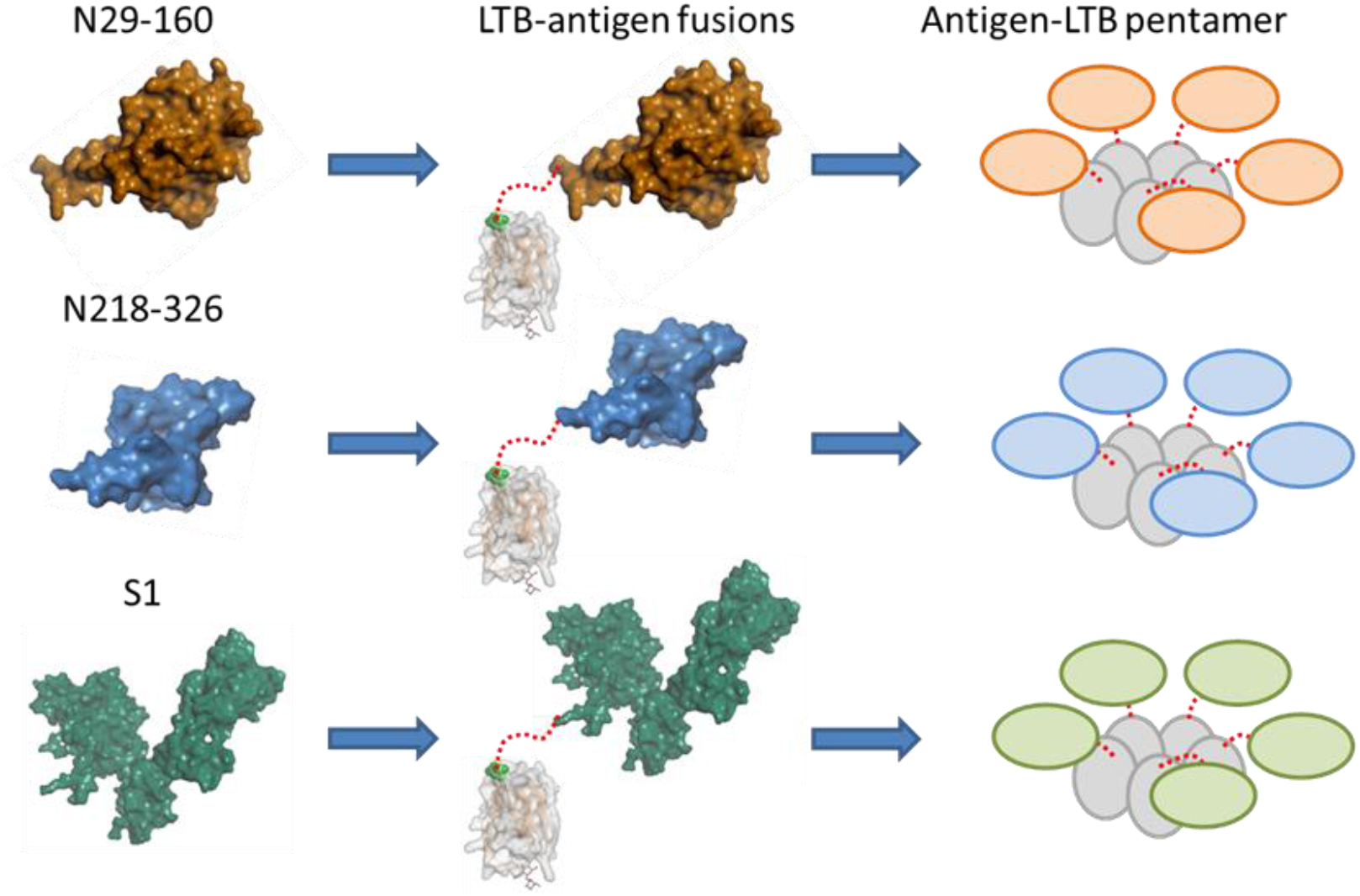
3D models of the selected antigens. The N-terminal and C-terminal segments of the nucleoprotein are shown in orange and blue-surface representation, respectively. The S1 segment is shown in green-surface representation. LTB is shown as a monomer in grey-surface representation and at its C-terminal residue, the attachment point for the antigens of each monomer, is presented in green, van Der Waals display. A schematic model of the pentamer composed of fused LTB (grey spheres)-antigen monomers is shown in the right column.

Since all fusion proteins contain the natural leader peptide from LTB, driving its secretion to the periplasm, the proteins were collected from the bacterial medium following shaking for 24-48 h at 37 °C. Partial purification was performed by ultrafiltration to increase the protein concentration. Briefly, 1L of the collected bacterial medium was passed through a 0.2 μm microbiological membrane and then concentrated x 100 using 10 kDa Amicon Ultra centrifugal filters (Merck Millipore). Positive clones were further verified by GM1 enzyme-linked immunoassay (ELISA), as detailed below.

#### Enzyme-linked immunosorbent assay

Fusion proteins were quantified by GM1 ELISA. The wells of a white polystyrene 96-well microtitre plate (Bio-One CELLSTAR, Greiner) were coated overnight, at 4 °C, with 100 μl bovine brain GM1 ganglioside (1.0 μg/ml) (Sigma-Aldrich) in 0.1 M carbonate-bicarbonate buffer, pH 9.5. The plates were washed five times with PBS added with 0.05% (v/v) Tween 20 (PBST), blocked by the addition of 2% skim milk (Difco) in PBST (1 h, 37°C, gentle shaking 70 rpm), and then washed again. Then, 100 μl of the expressed polypeptides, control peptides or standards were added to the wells (1h, 37°C, gentle shaking 70 rpm). Commercial LTB (Sigma) 100-1000 ng/ml was used as a standard for the calibration curve. CAYE medium served as a blank and filtrate from the wild type *E. coli* served as negative control. After washing, 100 μl rabbit serum anti-cholera toxin (Sigma-Aldrich) (1:15000 in PBST) was added to all wells and incubated (1h, 37°C, gentle shaking 70 rpm). After extensive washing, 100 μl HRP-labelled goat anti-rabbit IgG (Abcam) (1:10000 in PBST) were added to all wells (1 h, 37 °C). After a final washing, 100 μl Immobilon Crescendo western HRP substrate (Merck Millipore) was added to the wells and luminescence (cd/m2) was measured by an Infinite M200 Pro ELISA Plate Reader (Tecan). The concentration of LTB-fused proteins was calculated from the linear standard curve of LTB following subtraction of the blank.

### Chicken vaccination

All animal experiments were carried out in accordance with the guidelines of the Israeli Ethics Committee (approval ID IL-18-1-1).

#### Preparation of bacteria for oral vaccination

*E. coli* MG1655 secreting LNN, LNC or LS1 fusion proteins as well as WT bacteria were stored on Luria Bertani (LB) agar plates in 4 °C. In order to verify stability of the plasmids carrying the virus genes, each of the four bacterial groups was resuscitated by spreading a colony on a new LB agar plate four days before vaccination. The bacteria were cultured at 37°C, on a LB substrate with ampicillin (100 μg / ml). Bacterial starters were prepared from the renewed cultures, by growing one colony from each clone in 4 ml liquid LB substrate in a 50 ml test tube (37 °C, 24 h) on a RotamixRM mixer. A sample (100 μl) from each starter was transferred to tubes with 25 ml CAYE substrate and grown (48 h, 37°C, on a RotamixRM mixer). Thereafter, cultures were spectrophotometrically equilibrated to ~5×10^9^ CFU/mL and administered fresh to the chick in 1mL dose. M41 commercial inactivated IBV vaccine (Phibro, Israel) was administered intramuscularly at a final volume of 0.5 ml (as a positive control).

#### Experiment 1A- Evaluation of humoral immune activation

Chickens received the first vaccine dose at 30 days of age and a boost dose at the age of 44 days. Oral administrations were applied by pipetting 1ml partially purified protein (50 μg; n=5), prepared as described above, or cultured bacteria (1×10^9^ cells/ml; n=7) into the chicken’s oral pharynx.

The experimental design included five groups as specified in Table 2.

**Table 2.** Experiment 1 chicken groups.

| Group | N | Vaccine |
| --- | --- | --- |
| 1 | 7 | WT *E. coli*<sup>*</sup> |
| 2 | 7 | Mix of *E. coli* expressing subunit proteins LNC and LNN<sup>#</sup> |
| 3 | 7 | *E. coli* expressing subunit protein LS1<sup>†</sup> |
| 4 | 7 | *E. coli* expressing subunit proteins LS1, LNC and LNN |
| 5 | 5 | Partially purified LS1 protein<sup>‡</sup> |

<sup>\*</sup>WT: Wild type - Negative control *E.coli* that contain no plasmid
<sup>#</sup>LNC: C-terminus of IBV N protein fused to LTB, LNN - N-terminus of IBV N protein
fused to LTB
<sup>†</sup>LS1- Spike 1 protein fused to LTB
<sup>‡</sup>Protein collected from the medium

Fourteen days after the boost dose, chickens were bled and anti-IBV protein IgY levels were determined.

#### Experiment 1B- Evaluation of cellular immune activation with LTB-N sub units

Chickens (n=2 in each group) were vaccinated orally (as specified previously) or by subcutaneous injection (50 μg per chicken) of LNN or LNC. Fourteen days after the boost dose, chickens were sacrificed and spleens were harvested and evaluated for activation of cellular-mediated immunity.

#### Experiment 2 - Challenge test

Specific-pathogen-free, 1-day-old Leghorn chicks (n=63) were marked by wing tags and randomly separated to seven isolation units, under positive air pressure and artificial daytime lighting. The chicks received drinking water and commercial ration *ad lib*. Each bird in groups 1-6 was vaccinated intra-crop, 3 times, at 15, 36 and 45 days of age, with 1 ml of a CAYE bacterial substrate as specified in Table 3. The mixed bacterial substrates delivered to groups 5 and 6 were prepared in a 1:1 or 1:1:1 ratio, respectively, and a total of 1 ml of the mixture was administered. Chickens in group 7 were subcutaneously vaccinated with 0.5 ml commercial inactivated IBV vaccine (Virasin 227, Phibro, Israel). Following vaccination, chickens were inspected daily for clinical signs or mortality.

**Table 3.** Experiment 2 chicken groups.

| Group | N | Vaccine | Challenge |
| --- | --- | --- | --- |
| 1 | 3 | Caye broth (negative control) | - |
| 2 | 7 | Caye broth (positive control) | + |
| 3 | 9 | WT *E. coli*<sup>*</sup> | + |
| 4 | 7 | *E. coli* expressing subunit proteins LS1<sup>&</sup> | + |
| 5 | 13 | Mix of *E. coli* expressing subunit proteins LNC and LNN<sup>†</sup> | + |
| 6 | 13 | Mix of *E. coli* expressing subunit proteins LS1, LNC and LNN | + |
| 7 | 11 | Commercial inactivated M41 | + |

<sup>\*</sup>WT: Wild type - Negative control *E.coli* that contains no plasmid
<sup>&</sup>LS1: Spike 1 protein fused to LTB
<sup>†</sup>LNC: N-terminal part of IBV N protein fused to LTB, LNN: C-terminal part of IBV N protein fused to LTB

Fifteen days after the last boost vaccination, the chickens in groups 2-7 were challenged with the M41 strain of IBV at a viral titre of 10^7.5^ EID_50_/ml. More specifically, 200 μl challenge virus was divided to four portions of 50 μl each, which were administered into both eyes and both nostrils. Following challenge, the chickens were inspected daily for clinical signs or mortality. On the day of challenge and 3, 6 and 10 days post-challenge, choanal and cloacal swabs were collected from all chickens, to measure IBV shedding by q-PCR. Ten days post-challenge, the chickens were euthanized by CO_2_.

#### Determination of antibody levels

Serum was analysed for the presence of specific antibodies against the target IBV proteins, using a commercial IBV ELISA kit (Biocheck), according to the manufacturer’s instructions.

### Cell-mediated immunity evaluation

Two chickens randomly selected from each group in experiment 1 which were orally vaccinated with live bacteria, were sacrificed, and spleens were dissociated using a gentle MACS™ dissociator (Miltenyi Biotec) and then sieved through a 70 μm mesh to obtain a single-cell suspension. Splenocytes were suspended in RPMI 1640 supplemented with 2% foetal bovine serum, 2 mM L-glutamine, penicillin (100 U/ml) and streptomycin (10 ng/ml), (Biological Industries, Beit HaEmek, Israel). Cells (3 × 10^6^/well) were seeded in 24-well culture plates and incubated for 48 h at 37°C with lipopolysaccharide (LPS) (5μg/ml) or concanavalin-A (5μg/ml) as positive controls, PBS as negative control, or with formalin (0.3%)-inactivated IBV strain H120 (5×10^3.5^ EID_50_/ml; Vir 111, Biovac, Or Akiva, Israel). Splenocytes were transferred to fluorescence-activated cell sorting (FACS) tubes, washed twice by centrifugation (500 g, 4 °C) and suspended in cooled FACS buffer (PBS with 0.1% BSA). Samples were incubated (4 °C, 45 min) with antibodies against CD45 (APC), CD3 (biotin), CD4 (PE), and CD8 (FITC), all of which were purchased from Southern Biotec (Birmingham, AL, USA). Streptavidin-PE/Cy7 was purchased from Biolegend (San Diego, CA, USA). Samples were then washed twice and suspended in PBS. Flow cytometry was carried out using a FACSAria (Becton–Dickinson) and results were analysed with FCS express 4 software (De Novo Software). Activation was defined as a significant increase of the percentage of CD8^+^ cells in the CD3^+^ cell population.

### Determination of virus concentration using quantitative RT-PCR

Viral RNA from all swab samples collected in Experiment 2 was extracted using a RNA easy Mini Kit (QIAGEN), as per the manufacturer’s instructions. Primers and probe of N gene were as previously described [26]. Each PCR reaction included 5 μL RNA, 12.5 μL of 2X RT-PCR Buffer, 1 μL of each 10 μM primer, 1 μL of 3 μM TaqMan probe, 1 μL of 25X RT-PCR Enzyme Mix and 3.5 μL H_2_O, in a total reaction volume of 25 μL. The RT-PCR was performed using AgPath-ID™ One-Step RT-PCR Kit (Thermofisher). The reaction protocol was as follows: one cycle at 45 °C for 10 min, one cycle at 95 °C for 1 min and 40 cycles (95 °C for 15 sec and then 60°C for 45 sec) for amplification. A standard curve was prepared by serially diluting IBV M41 strain (10^6.5^ EID_50_/ml) in PBS and subjecting the RNA extracted from each dilution to RT-PCR, as described above. The standard curve results were used to determine virus concentrations in swab samples. Real-time RT-PCR cycle threshold (CT) >36 was considered negative.

### Statistical analysis

The data are expressed as means ± standard deviation (SD). Significant differences between two groups were determined by the unpaired two-tailed Student’s t-test. For multiple mean comparisons, 1-way analysis of variance (ANOVA) with Tukey HSD post-hoc test was performed. p≤0.05 was considered statistically significant. Chi-squared and Fisher’s exact test were carried out to assess intergroup differences in the percentage of birds that shed IBV.

## Results

### Antigen selection and fusion to carrier protein

The entire S1 segment and two separate segments of the nucleoprotein (NN: amino acids 29-160 and NC: amino acids 218-326) were selected based on analysis of the available high-resolution structures of both S1 and N segments of homologue proteins (Figure 1).

### Expression and secretion of the recombinant proteins

The presence of the expressed proteins in the bacterial growth medium was verified by Western blot analysis. Protein bands of molecular weights of 26 kDa, 29 kDa and 72 kDa were in concordance with the predicted molecular weight of LNC, LNN and LS1, respectively (data not shown). LTB secretion was verified by ELISA, which demonstrated that all fusion proteins were secreted to the medium in a time-dependent manner. Concentrations measured in the medium following 48 h of bacterial growth and protein expression were 800 ng/ml, 350 ng/ml and 200 ng/ml for LNC, LNN, and LS1, respectively (Figure 2).

**Figure 2.**
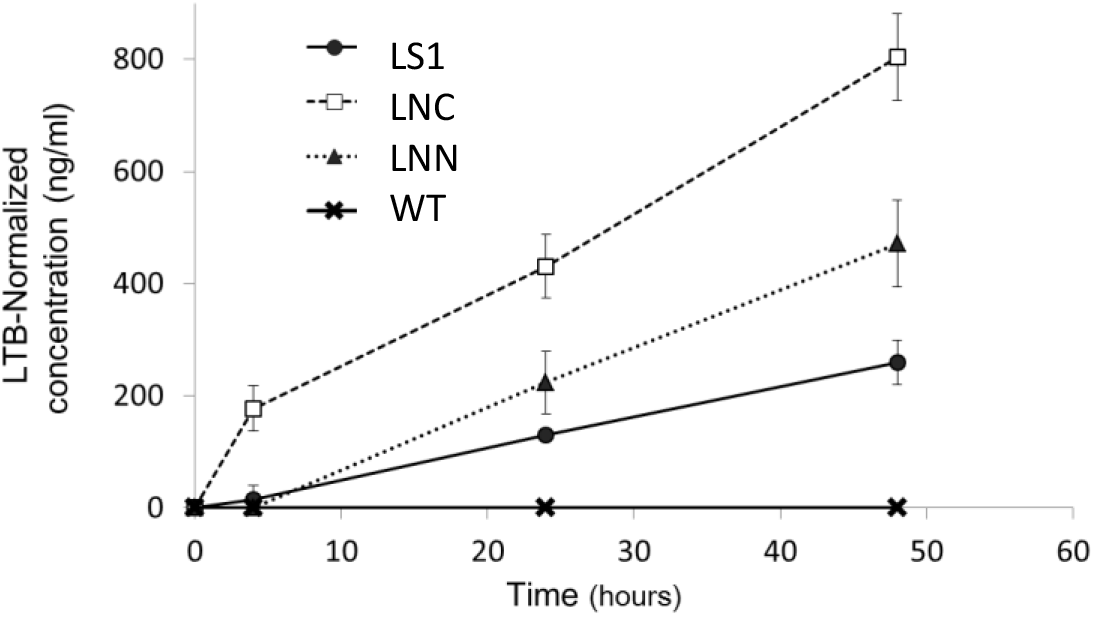
Kinetics of fusion protein secretion to bacterial growth medium. *E. coli* MG1655 wild type (WT) as negative control or clones expressing LS1, LNC or LNN, were cultured. Growth medium (100 μl) was collected after 0, 6, 24 and 48 h. concentrations were determined by ELISA. Proteins concentrations at each time point were calculated based on linear standard curve of serially diluted commercial LTB. The graph presents the average ± standard deviation of two independent experiments, each performed in triplicates.

### Immune response following vaccination with LNN, LNC and LS1 fusion proteins

#### Antibody response

Orally administered vaccines containing chimeric LNN, LNC and LS1-secreting bacteria, or partially purified and concentrated LS1, induced antibody levels that exceeded the kit’s positive cut-off (Figure 3). Vaccination with the mix of bacteria secreting LNN, LNC and LS1 domains resulted in higher antibody levels as compared to vaccination separately with the two LNs or LS1 domain. Chicks orally vaccinated with the partially purified and concentrated LS1 exhibited slightly higher but comparable antibody levels as the group receiving the LS1-LNN-LNC vaccine mix.

**Figure 3.**
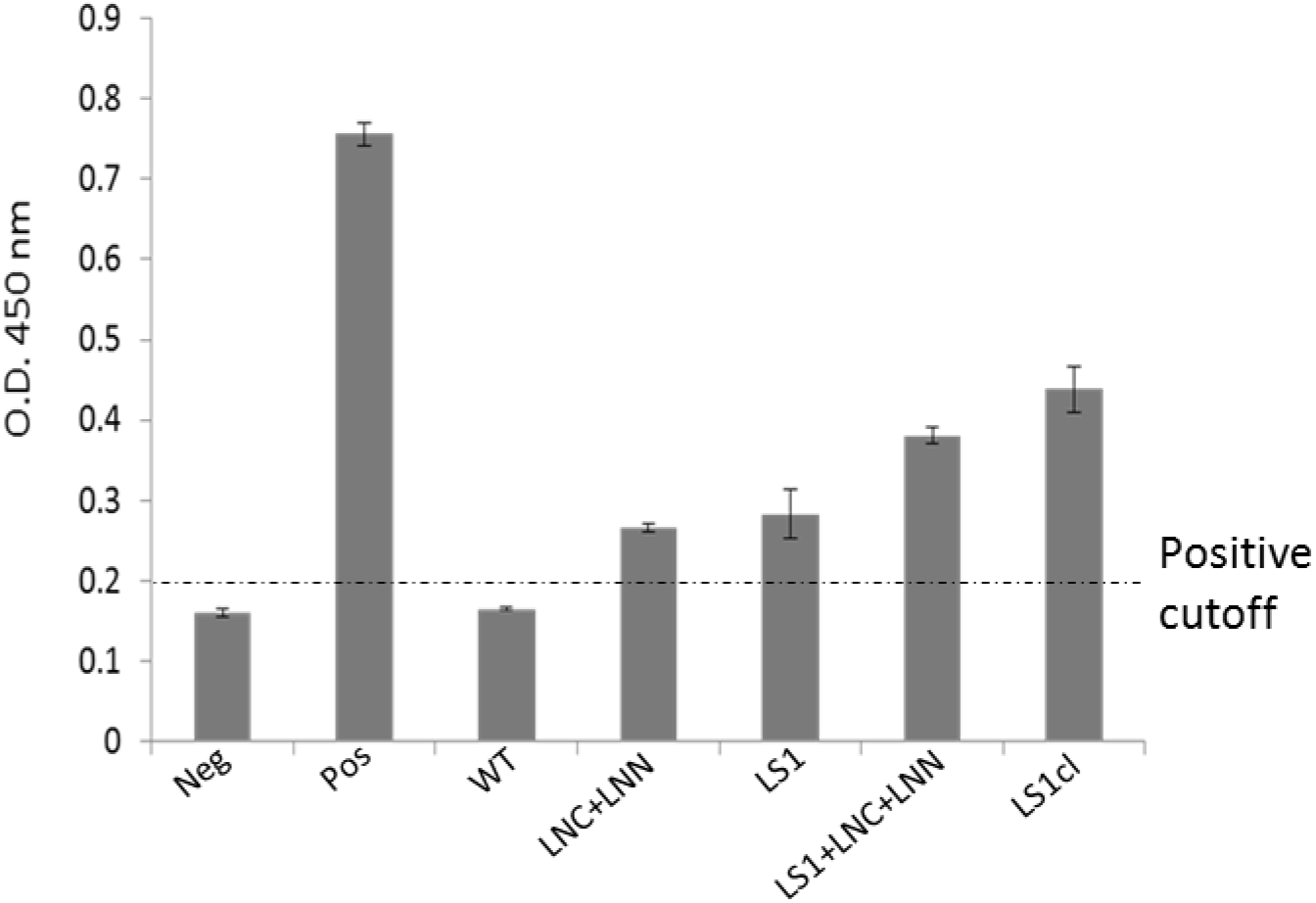
Level of anti-IBV IgY antibodies following oral vaccination. A commercial IBV-ELISA kit was used to determine antibody levels in chicks vaccinated with live bacteria expressing LTB-fused infectious bronchitis virus (IBV) LS1, LNC+LNN or LS1+LNC+LNN polypeptides. Neg and Pos represent the commercially provided kits negative and positive controls, respectively. LS1cl chickens were vaccinated with partially purified LS1 polypeptide. Each bar represents the average ± standard deviation of antibody levels. Sera samples from each vaccination group (n=5-7 chickens per group) were pooled and tested in triplicates. Positive cut-off (dashed line) signifies efficacious vaccination or previous exposure as stated by the manufacturer.

#### Cellular immune response

When exposed to inactivated IBV, splenocytes from chickens orally vaccinated with bacteria secreting LNN or LNC fusion proteins, showed a significantly higher percentage of CD8^+^ cells among the CD3^+^ population as compared to splenocytes isolated from chickens vaccinated with WT or LTB-secreting *E. coli* (Figure 4A). Moreover, oral administration of the LNN fusion protein yielded significantly higher %CD8 levels as compared to vaccination with the same protein administered by injection (Figure 4B).

**Figure 4.**
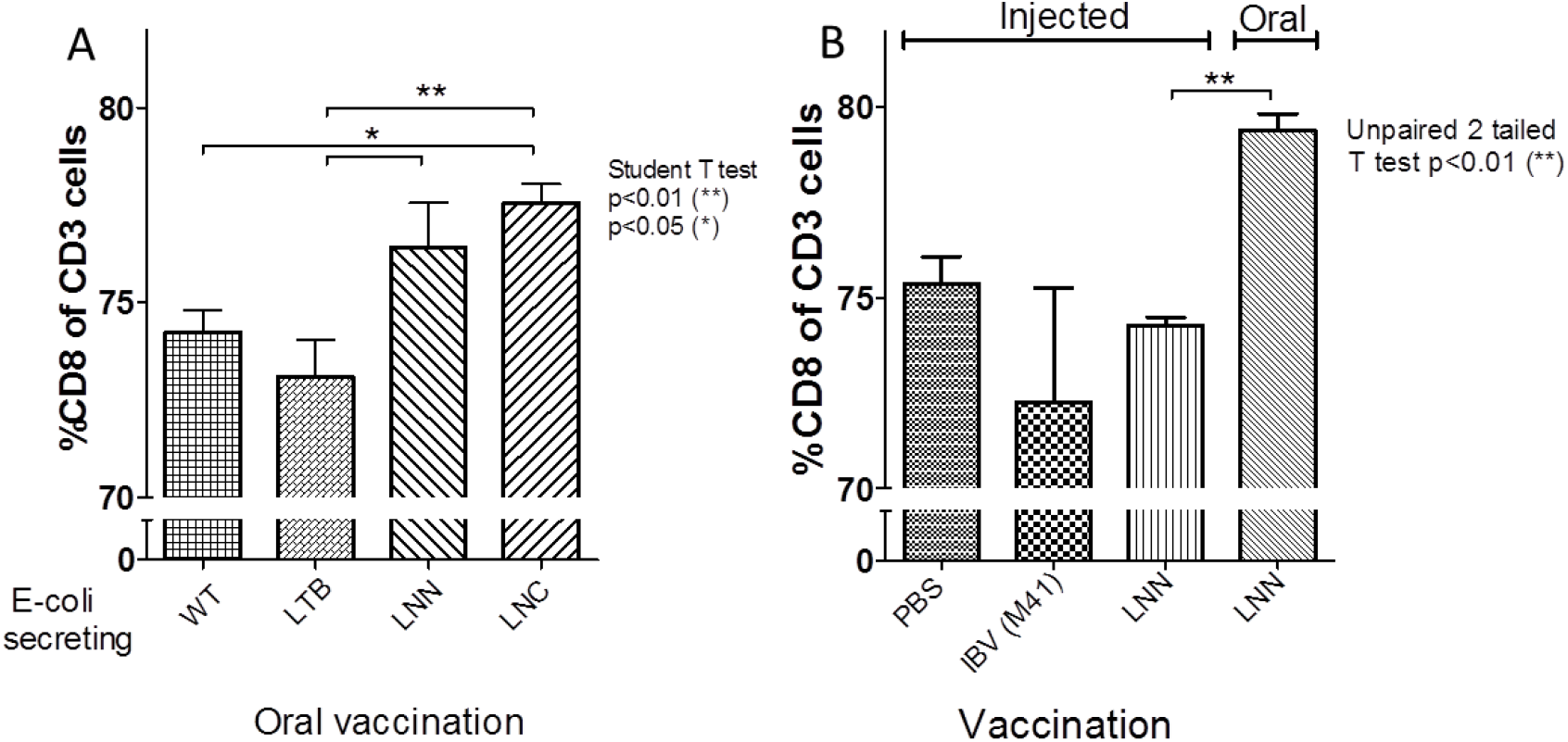
Nucleocapsid (N)-induced cell-mediated immunity. Splenocytes collected from chickens orally vaccinated with *E. coli* secreting various IBV components were incubated with formalin-inactivated IBV strain H120 and then subjected to fluorescence-activated cell sorting (FACS) to determine percentages of CD8+ cells out of total CD3 T cell population. A. Chickens were orally vaccinated with control wild type *E. coli* (WT), *E.coli* expressing LTB, LNN fusion, or LNC fusion protein. B. Chickens were injected with PBS, inactivated IBV (strain M41) or LNN, or orally administered LNN. Bars marked with an asterisk indicate significant differences in percentages between the different groups *p<0.05, **p<0.01.

### Challenge study

#### Virus shedding

The concentrations of the virus shed over time from the choana and the trachea following challenge with live, virulent IBV are presented in Fig. 5. Virus concentrations in the group vaccinated with *E. coli* expressing LS1+LNC+LNN decreased significantly in the choana from day 0 to day 10 post-challenge and were significantly lower than in all other challenged groups. More specifically, on day 3 post-challenge, virus shedding from the choana was about 1.7 Log_10_ EID_50_/ml in the birds vaccinated with LS1+LNC+LNN, and about 5 Log_10_ EID_50_/ml in all other treatment groups. On days 6 and 10 post-challenge, virus titres for this group dropped below 1 Log_10_ EID_50_/mL. In all other challenged groups, titres of shed virus in the choana decreased only slightly by day 6. On day 10 post-challenge, only chicks vaccinated with LS1 or LS1+LNC+LNN showed shedding similar to that of the non-challenged negative control group. In all other groups, titres dropped below 2 Log_10_ EID_50_/ml, with high variability among the birds of the each group. Due to this large variance within groups, statistical differences between groups were small. Nevertheless, titres of viruses shed on day 6 by chicks vaccinated with LS1+LNC+LNN were significantly lower than those found in chick groups vaccinated with, IBV (M41) or LNC+LNN control groups.

**Figure 5.**
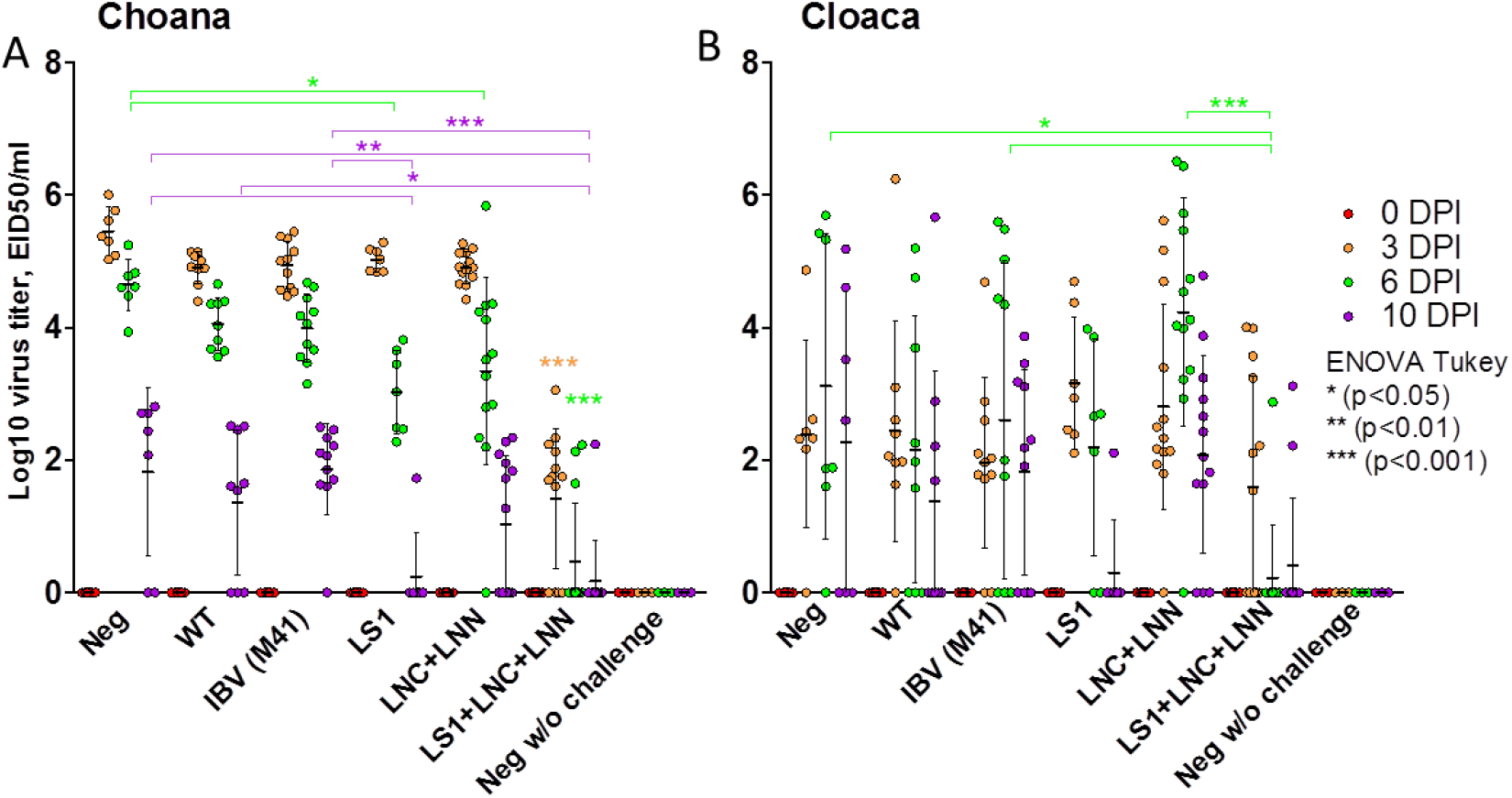
Levels of IBV shedding in birds following IBV challenge. Chickens were challenged with live virulent IBV (strain M41) 15 days after administration of a last booster dose of either a subcutaneously injected commercial inactivated IBV (strain M41) vaccine, or orally administered bacterial Caye broth growth medium (neg), wild type *E. coli* (WT), or *E. coli* strains secreting the LTB-fused polypeptides: LS1, 1:1 ratio of LNC and LNN (LNC+LNN) or 1:1:1 ratio of LS1, LNC and LNN (LS1+LNC+LNN). Graphs show the IBV titres shed by chickens, as determined by qRT-PCR performed on swab samples collected from either the choana **(A)** or the cloaca **(B)**, before challenge (0), and 3, 6 and 10 days post-infection (DPI). The differences between groups at 3, 6 and 10 DPI were analysed by one-way-ANOVA with Tukey HSD post-hoc. Statistical analyses for groups at 3, 6 and 10 DPI are presented in orange, purple and green respectively.

A time-course analysis of viral shedding from the choana or cloaca following challenge (day 0), showed that all vaccinated birds shed IBV 3 days after the challenge, apart from the group vaccinated with LS1+LNC+LNN (Table 3, group 6), in which only 85% were shedders (Figure 6). By 6 days post-challenge, only 23% of the LS1+LNC+LNN-vaccinated chickens were IBV shedders, while all bird in all other groups remained positive. On day 10 post-challenge, only 14% and 23% of the chickens vaccinated with LS1 or LS1+LNC+LNN, respectively, were shedders, a significantly lower percentage than the percentages found in the other test and control groups. In contrast, all of the LNC+LNN-vaccinated chicks continued to shed IBV until day 10 post-challenge.

**Figure 6.**
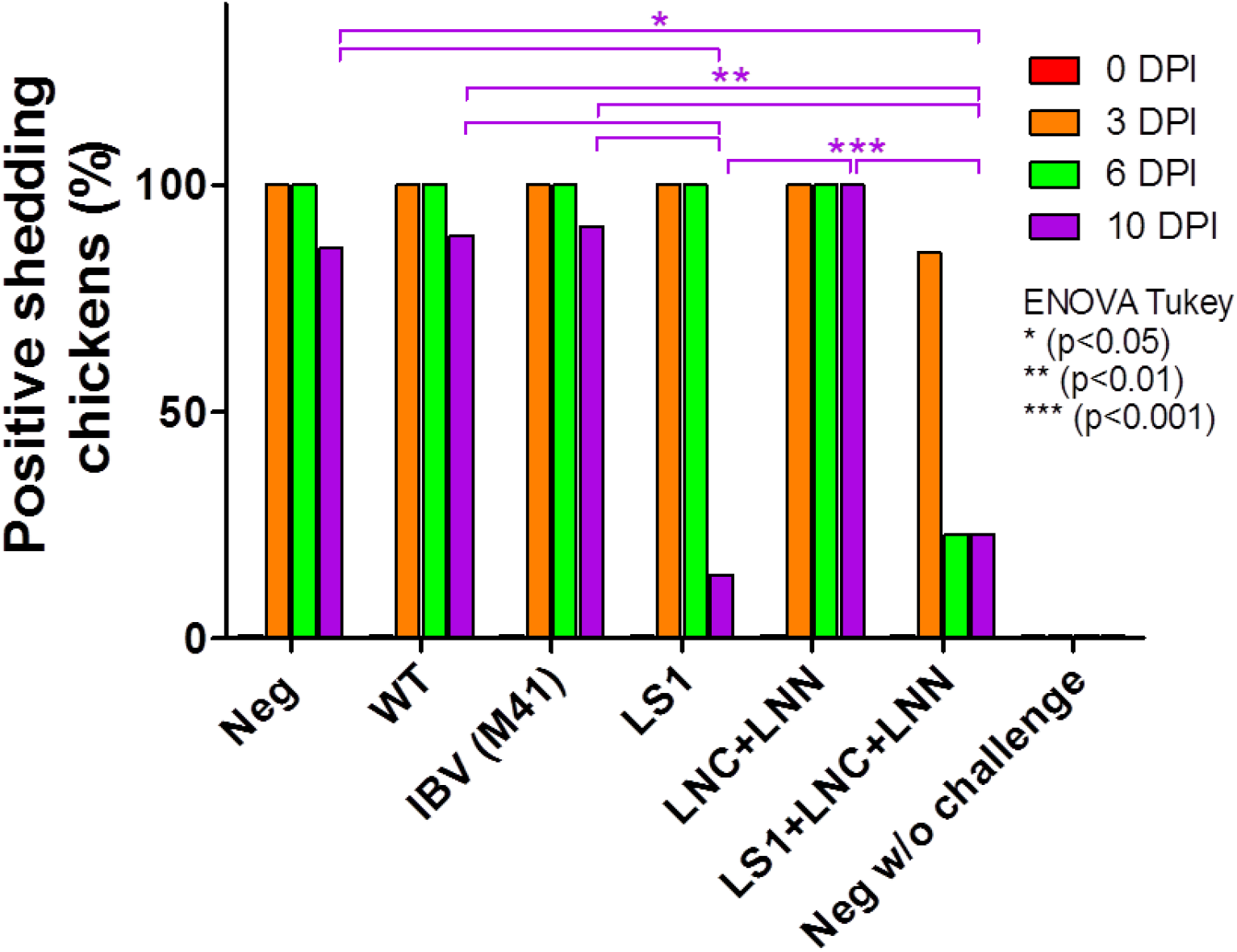
Percentage of birds that shed IBV from either the choana or the cloaca following IBV challenge. Chickens were challenged with live virulent IBV (strain M41) 15 days after administration of a last booster dose of either a subcutaneously injected inactivated IBV (strain M41) vaccine, or orally administered bacterial Caye broth growth medium (neg), wild type *E. coli* (WT), or *E. coli* strains secreting the LTB-fused polypeptides: LS1, 1:1 ratio of LNC and LNN (LNC+LNN) or 1:1:1 ratio of LS1, LNC and LNN (LS1+LNC+LNN). Graphs show the percent of shedding chickens as determined by qRT-PCR performed on swab samples taken from either the choana or the cloaca, before challenge (0), and 3, 6 and 10 days post-infection (DPI). The differences between groups at 10 DPI were analysed by Fisher’s exact test.

## Discussion

This work described the design and evaluation of an oral anti-IBV vaccine comprised of 3 LTB-fused IBV polypeptides delivered using a live bacterial platform. Vaccine design optimized for LTB fusion and epitope presentation using high-resolution structural models, resulted in proteins that were expressed and secreted by *E. coli*. When orally delivered to chickens, the IBV polypeptide-expressing bacteria induced production of virus-specific IgY antibodies. Orally administered LTB-NN was associated with splenocyte immune responses following exposure to an inactivated IBV that exceeded the responsiveness of splenocytes isolated from chickens subcutaneously vaccinated with the same polypeptide. In addition, significantly less viral shedding was measured in chickens orally immunized with the LTB-fused IBV peptide mix or with LS1 as compared to all other test groups, including those receiving the commercial inactivated vaccine. In the group vaccinated by the LTB-fused IBV peptide mix, a significantly shorter shedding period was shown as compared to all other groups, including the group vaccinated with LS1 alone.

Mucosal vaccination confers benefits over other routes of administration, including strong mucosal immunity, which provides the first barrier against pathogens entering via the mucosa. Live attenuated viruses are widely used for mucosal vaccination, including Newcastle disease virus [27], IBV [28] and avian reovirus [29]. Yet, the attenuation process is long and costly and some safety concerns are raised, such as the possibility of virus reversion to a virulent state in immunocompromised hosts [30]. Protein-based vaccines are intrinsically safer than whole pathogen-based vaccines and are faster to adjust. Yet, despite extensive efforts toward development of protein-based mucosal vaccines against various infectious diseases, there are currently no approved oral or intranasal protein vaccines. Integration of LTB in protein vaccines has been shown to enhance immune responses. In addition, live bacterial cells as delivery vehicles for recombinant antigen have been suggested to serve as effective adjuvants due to their potent activation of the innate immune responses [18, 31]. For example, a recombinant subunit vaccine containing the R repeat region of P97 adhesin of *M. hyopneumoniae* fused to LTB and expressed in *E. coli*, produced high levels of systemic and mucosal antibodies in intranasally or intramuscularly inoculated BALB/c mice [32]. Similarly, LTB-fused recombinant C-terminal fragments of botulinum neurotoxins serotypes C and D, produced in *E. coli*, induced high levels of neutralizing antibodies [33]. The same recombinant vaccine induced a strong immunogenic response in cattle as well [34]. Nandre et al. report on a significant reduction in egg and internal organ contamination by virulent *Salmonella* enteritidis (SE) in laying hens orally inoculated with live attenuated LTB-secreting *Salmonella* enteritidis (SE-LTB) as compared to those receiving a commercial anti-SE inactivated vaccine [35]. In line with these studies, the current work demonstrated the safety and potency of *E. coli* as a vaccine delivery platform, as well as the additive effect of LTB integration in vaccine design.

The presented vaccine was designed to simultaneously target several epitopes, with the goal of increasing humoral, mucosal and cellular immunity and providing for more robust control and prevention of IBV disease. The research focused on the S and N proteins, in light of evidence of the role of both S1 and N genes in the induction of immune responses against IBV [36–38]. Specifically, Meir et al. [39] reported on both humoral and cell-mediated immune responses following vaccination of chickens with an anti-IB V eye drop vaccine comprised of recombinant S1 or N expressed in *E. coli*. The selected peptides correspond to regions of both the S1 and N viral proteins that provide maximal coverage of protective epitopes. Integration of the highly conserved N protein may contribute to group-common immunity against IBV variant viruses. These features were ensured by a thorough structural analysis of available high-resolution structures of the S1 and N homologue proteins in search of independent folding units, while avoiding destabilization of the native structure. Fusion of these antigen sequences to LTB via a linker segment allowed both native LTB pentameric complex formation and the native fold of each of the antigens S1 and N. The construct design was aided by use of 3D computer modelling available for IBV proteins S and N. For the S1 protein sequence analysis, the cryo-electron microscopy structure of infectious bronchitis coronavirus spike protein was used [40]. This structure is 96.5 identical to the S1 sequence from strain H120 and therefore served as the template for homology model of the S1 protein. The N protein had fragmented structures from similar sequences in the protein data bank spanning residues 22-160 and 218-333 [41, 42]. The N-terminal part (residues 22-160) of the high-resolution structures of the nucleoprotein is 90.3% identical to the correlated segment from H120 strain used in this study and the C-terminal segment (residues 218-333) of the high-resolution structures of the nucleoprotein is 93.6% identical to the correlated segment from H120 strain used in this study.

Poultry rearing, one of the largest livestock enterprises, is highly vulnerable to viral outbreaks, which can have devastating implications on local and national economies, as well as on food security. Thus, breeders are continuously seeking potent and cost-effective vaccination strategies to control disease. Oral inoculation is a user-friendly mass-vaccination strategy. Given that IBV is primarily transmitted via respiratory droplets, effective mucosal immunity might improve protection against nasal and/or oral virus entry. Moreover, mucosal immunity decreases viral shedding, as described in the current study. In addition, this route promises to overcome significant constraints related to vaccine administration, including the avoidance of stress due to chicken herding and injection. The rationale behind the presented vaccine design may be the basis for further development of affordable, rapidly adaptable mucosal non-live vaccines.

## Acknowledgements

The authors would like to thank Prof. Dan Heller and Dr. Amnon Michael for their highly valued contribution in the weaving and orchestration of the Center for Vaccine Development of Viral Diseases of Poultry.

